# Multiomic Integration Reveals Transposable Element–Embedded Regulatory Variants Underlying Brain Aging

**DOI:** 10.64898/2026.07.28.740948

**Authors:** Sreejato Chatterjee, Shuangshuang Feng, Dean Zhang, Hongbo Liu

**Affiliations:** Department of Biology, University of Rochester, Rochester, NY, USA; Department of Biomedical Genetics, University of Rochester Medical Center, Rochester, NY, USA; University of Rochester Aging Institute, University of Rochester Medical Center, Rochester, NY, USA; Wilmot Cancer Institute, University of Rochester Medical Center, Rochester, NY, USA

**Keywords:** genome-wide association study, brain aging, transposable elements, epigenomics, chromatin accessibility, ENCODE cCREs, single-nucleus ATAC-seq, ChromHMM, enhancer–promoter interactions, variant-to-gene prioritization, functional genomics

## Abstract

Genome-wide association studies (GWAS) have identified thousands of variants associated with brain aging, most in noncoding regions whose regulatory effects remain poorly understood. Transposable elements (TEs) constitute a major, understudied component of brain regulatory DNA. To interpret noncoding brain-aging variants through TE-derived regulatory architecture, we integrated brain-aging GWAS variants with TE annotations, ENCODE candidate *cis*-regulatory elements (cCREs), single-nucleus ATAC-seq accessibility maps, HOMER motif analysis, experimental enhancer–promoter interaction maps, and multiomic variant-to-gene resources. Of 6,323 genome-wide-significant variants, 633 (10.0%) localized within 367 TE-derived cCREs. These elements were predominantly enhancer-like and frequently carried transcription factor–associated annotation. Alu-derived elements formed the largest family component, followed by L1, L2, MIR and ERVL-MaLR elements. Single-nucleus accessibility maps resolved these variants across brain cell types, including a TcMar-Tigger–associated signal in oligodendrocytes, and motif analysis of SINE-associated cCREs identified reproducible nuclear-receptor, immune (PU.1:IRF8) and chromatin-regulatory (RFX-family) signatures. Because this localization reflects correlated variants within a limited number of independent loci rather than independent enrichment signals, we prioritized candidate regulatory variants and their target genes using the Brain Aging Genetic ScoreCard, which integrates 27 orthogonal evidence layers across independent, LD-defined loci and identified 25 loci with TE-derived regulatory support. The chromosome 17q21.31 *MAPT* locus emerged as the top candidate genome-wide, where a MIR-derived enhancer shows convergent evidence linking it to *MAPT*, a prioritization robust to locus-size effects. Overall, our study provides an integrative framework for interpreting noncoding GWAS variation through TE-derived regulatory architecture.

## 1 Introduction

Human brain aging is a multifaceted and heterogeneous biological process, strongly shaped by epigenetic regulation, which maintains transcriptional stability and genome integrity in long-lived neuronal cells. Unlike rapidly dividing tissues, brain cells must preserve gene expression programs over decades, relying on tightly controlled chromatin states to prevent aberrant transcription and genomic instability. Disruption of these regulatory mechanisms has been implicated in age-related cognitive decline and neurodegenerative disease, highlighting the importance of genetic control in maintaining neuronal function over the lifespan [1, 15].

Genome-wide association studies (GWAS) have identified many loci associated with brain-aging-related phenotypes, including cognitive function, brain structure, and neurodegenerative disease risk. However, most of these variants lie in noncoding regions of the genome, where their biological effects cannot be inferred from protein sequence changes alone [14]. Instead, noncoding variants are thought to act through gene regulatory mechanisms, influencing when, where, and how genes are expressed. Interpreting these variants therefore requires placing them within an epigenetic and regulatory context. Frameworks integrating enhancer activity, chromatin contact, molecular quantitative trait loci (QTLs), and functional genomic evidence, including Activity-by-Contact (ABC) models and variant-to-gene prioritization resources, have therefore become central for linking noncoding variants to candidate target genes [4, 13, 5, 12].

This challenge is particularly important in the brain, where gene regulation is highly cell-type-specific. Distinct populations of neurons and glial cells exhibit different chromatin accessibility landscapes and regulatory programs, meaning that the functional impact of a given variant depends on the cellular context in which it resides [21]. ENCODE candidate *cis*-regulatory elements (cCREs) classify genomic regions into functional categories such as promoter-like, enhancer-like, and transcription factor-associated elements based on epigenomic signatures [19]. Complementary to this, Single-nucleus ATAC-seq (snATAC-seq) enables mapping of accessible chromatin across individual brain cell types, providing a framework for identifying where regulatory DNA is active. Integrating GWAS variants with these annotations allows noncoding loci to be interpreted in terms of both their cellular context and regulatory function.

Transposable elements (TEs) represent a major component of this regulatory landscape. Rather than acting solely as genomic parasites, TEs are frequent targets of epigenetic repression and can serve as sources of regulatory DNA, including transcription factor binding sites and enhancer elements [22]. More than 40% of the human genome is derived from TE sequences, and a substantial fraction of regulatory elements show TE origin [2]. Recent large-scale annotation efforts have demonstrated that approximately one-quarter of human candidate *cis*-regulatory elements (cCREs) are derived from transposable elements, yet the function of these TE-derived cCREs in brain aging is largely unknown. In the brain, TE activity is tightly controlled through epigenetic mechanisms, and dysregulation of these elements has been linked to aging and neurodegenerative processes [15]. These properties make TE-associated sequences plausible substrates through which noncoding genetic variation may influence gene regulation in aging brain tissue.

Among TE classes, SINE elements, including Alu and MIR elements, are of particular interest in primate brains. Alu elements are primate-specific, highly abundant, and enriched for transcription factor binding motifs relevant to neuronal function. Prior work has shown that Alu-derived sequences can function as enhancers and are frequently embedded within accessible chromatin in neural tissues [16, 20]. MIR elements are evolutionarily ancient, non-autonomous tRNA-derived retrotransposons approximately 260 nucleotides (nt) in length that harbor a conserved CORE-SINE motif. They are widely distributed across mammalian genomes and constitute approximately 2.54% of the human genome. However, it remains unclear whether noncoding brain-aging GWAS variants within SINE elements are preferentially enriched within functionally validated TE-derived regulatory elements, and whether any observed enrichment reflects properties of specific TE families or broader regulatory architectures shared across the genome.

In this study, we investigated whether brain-aging GWAS variants are preferentially localized within TE-derived regulatory elements and how these loci contribute to regulatory architecture across brain cell types. Specifically, we addressed two related questions: (1) Are brain-aging GWAS variants enriched within TE-derived candidate *cis*-regulatory elements (cCREs), and if so, which TE classes and families drive this enrichment? and (2) through which regulatory mechanisms, chromatin contexts, and downstream regulatory interactions might these variants exert their effects? To address these questions, we integrated brain-aging GWAS variants with RepeatMasker TE annotations, the ENCODE TE-derived cCRE atlas, single-nucleus ATAC-seq chromatin accessibility maps across major brain cell types, and enhancer-promoter interaction maps. We first evaluated whether TE-derived regulatory elements were enriched among independent BAG signals, and then characterized the regulatory composition of TE-overlapping loci through cCRE classification, transcription factor motif enrichment, and enhancer-promoter interaction analyses. Through this integrative framework, we distinguish genomic localization from validated regulatory activity, downstream regulatory connectivity, and heritability contribution, providing a multi-layered assessment of how transposable-element-derived regulatory sequences shape the interpretation of noncoding variation in brain aging.

## 2 Methods

### 2.1 Data Sources & Preprocessing

Brain aging-related genome-wide association summary statistics were obtained from the large-scale genome-wide association study of brain age gap (BAG) by Jawinski et al. [7]. BAG is defined as the difference between MRI-predicted brain age and chronological age and is an established biomarker of accelerated brain aging associated with cognitive decline, neurodegenerative disease risk, and age-related structural brain changes. We analyzed the European-ancestry meta-analysis of the combined gray- and white-matter BAG phenotype, comprising 54,890 individuals from the UK Biobank discovery cohort (*n* = 32,634), UK Biobank European replication cohort (*n* = 20,423), and LIFE-Adult cohort (*n* = 1,833). The original study performed genotype quality control, imputation, association testing, and meta-analysis, releasing summary statistics for 9,628,868 autosomal variants in the GRCh37/hg19 reference genome. The present study did not repeat association testing, but instead performed downstream functional annotation and regulatory interpretation of the published GWAS signals. The original summary statistics are available from Zenodo (DOI: 10.5281/zenodo.14826943).

Genome-wide significant variants were defined using the conventional threshold of *P* < 5 × 10^−8^, yielding 6,479 significant variants. To ensure reliable representation in the linkage disequilibrium (LD) reference panel, analyses were restricted to common variants (minor allele frequency ≥ 0.05). Variant coordinates were converted from GRCh37/hg19 to GRCh38/hg38 using the UCSC liftOver tool and the hg19ToHg38 chain file, after which 6,323 genome-wide significant variants remained for downstream analysis. These variants were subsequently used for functional annotation with RepeatMasker, ENCODE 4 candidate cis-regulatory elements (cCREs), Roadmap ChromHMM states, Activity-by-Contact (ABC) enhancer–gene predictions, single-nucleus ATAC-seq data, and other regulatory datasets, all of which are available in the GRCh38 genome build.

### 2.2 Annotation of Transposable Elements

Genome-wide annotations of transposable elements were obtained from RepeatMasker for the hg38 reference genome. RepeatMasker classifies repetitive elements hierarchically into TE classes (e.g., SINE, LINE, LTR, DNA transposons) and families (e.g., Alu, MIR, L1). GWAS variants were intersected with RepeatMasker annotations using BEDTools intersect. A variant was classified as TE-overlapping if it overlapped any annotated TE by at least one base pair. Variants overlapping multiple TE annotations were retained with all corresponding TE labels to allow class- and family-level analyses.

### 2.3 Annotation of Candidate *cis*-Regulatory Elements

To further classify regulatory DNA into functional categories, we obtained the latest released candidate *cis*-regulatory element (cCRE) annotations from the ENCODE 4 project for the hg38 reference genome [11]. These ENCODE cCREs were classified as putative regulatory DNA into functional categories based on chromatin accessibility and histone modification signatures, including promoter-like signatures (PLS), proximal enhancer-like signatures (pELS), distal enhancer-like signatures (dELS), CTCF-associated elements (CA-CTCF), H3K4me3-associated elements (CA-H3K4me3), transcription factor-associated elements (CA-TF), chromatin-accessible elements (CA), and transcription factor binding elements (TF). GWAS variants were intersected with ENCODE cCRE annotations using BEDTools intersect. Because some variants overlapped multiple cCRE annotations, each variant was assigned a single representative cCRE class using a priority scheme reflecting regulatory specificity: PLS > pELS > dELS > CA-CTCF > CA-H3K4me3 > CA-TF > CA > TF.

### 2.4 TE-Derived Regulatory Element Enrichment Analysis

TE-derived cCREs were defined as ENCODE cCREs with at least 50% of their region covered by a single RepeatMasker-annotated transposable element. This criterion was applied using BEDTools intersect (-f 0.50) between the full ENCODE 4 cCRE registry (2,348,854 elements) and genome-wide RepeatMasker TE annotations (hg38). In rare cases where a cCRE satisfied the ≥50% threshold with more than one TE annotation (0.09% of qualifying cCREs), the single TE with the largest overlap fraction was retained. This yielded 759,488 TE-derived cCREs genome-wide, classified using the same eight-category ENCODE 4 functional taxonomy described in Section 2.3 (PLS, pELS, dELS, CA-CTCF, CA-H3K4me3, CA-TF, CA, TF).

Brain-aging GWAS variants were intersected with this TE-derived cCRE set using BEDTools intersect. Enrichment was evaluated relative to a background set of common variants (MAF > 0.05) from the 1000 Genomes Phase 3 European reference panel, independently intersected with the same TE-derived cCRE annotation.

### 2.5 Enrichment Analyses of TE-associated GWAS variants in cCREs

Enrichment analyses were performed to determine whether TE-associated GWAS variants were preferentially localized within specific regulatory annotations. For cCRE analyses, contingency tables were constructed comparing the distribution of TE-overlapping versus non-TE variants across cCRE classes. Overall differences were evaluated using chi-square tests, and standardized residuals were examined to identify regulatory categories contributing most strongly to the observed association. For individual cCRE classes, Fisher’s exact tests were performed comparing each class against all remaining classes, and odds ratios with 95% confidence intervals were calculated.

To assess enrichment within accessible chromatin, contingency tables compared TE-overlapping and non-TE variants inside versus outside ATAC-seq peaks for each brain cell type. Fisher’s exact tests evaluated enrichment and odds ratios quantified effect sizes. Multiple hypothesis testing across cCRE classes, TE families, traits, and cell types was controlled using the Benjamini–Hochberg FDR procedure. Statistical significance was defined as FDR-adjusted *p* < 0.05.

### 2.6 TE-derived promoter activity analysis

To evaluate whether brain-aging GWAS variants were enriched near TE-derived transcription start sites (TE-TSSs), we used a recently published catalog of candidate TE-derived TSSs identified from ENCODE RAMPAGE, RNA-seq, and ChIP-seq datasets [22]. TE-TSS coordinates were converted into BED format using peak chromosome, start, and end positions. Because experimentally defined TSS peaks are narrow, we performed two analyses: (1) direct overlap with annotated TE-TSS intervals and (2) overlap with ±500 bp windows centered on TE-TSS regions. TE-TSS intervals were restricted to autosomes and merged across samples to avoid overcounting recurrent TSS calls across tissues. Brain-aging independent GWAS variants and background variants were intersected with TE-TSS intervals using BEDTools.

### 2.7 Roadmap ChromHMM brain-state annotation

To assess tissue-specific chromatin context, TE-derived cCRE-associated GWAS variants were integrated with Roadmap Epigenomics 15-state ChromHMM annotations [3]. We used hg38-lifted ChromHMM mnemonic BED files from six brain tissues (angular gyrus, caudate, cingulate gyrus, hippocampus, dorsolateral prefrontal cortex [DLPFC], and substantia nigra) and two non-brain comparator tissues (liver and kidney) to assess the tissue specificity of TE-derived regulatory enrichment. Liver and fetal kidney ChromHMM segmentations were obtained from the same Roadmap Epigenomics core-15-state model and lifted to hg38 using the same procedure applied to the brain tissue tracks. ChromHMM states included active promoter, transcription-associated, enhancer, Polycomb-repressed, heterochromatin, and quiescent annotations.

Brain-aging GWAS variants overlapping TE-derived cCREs were intersected with ChromHMM state annotations using BEDTools, using the TE-derived cCRE interval containing each overlapping variant as the query region. As a background comparison, all TE-derived cCREs genome-wide were intersected with the same ChromHMM tracks. For each tissue (brain and non-brain) and ChromHMM state, contingency tables compared the number of GWAS-associated TE-derived cCRE overlaps with that state against the corresponding number of background TE-derived cCRE overlaps. Fisher’s exact tests were used to calculate odds ratios and p-values, with Benjamini–Hochberg FDR correction applied across all tissue-by-state comparisons.

ChromHMM states were also grouped into major functional categories including promoter-associated (TssA, TssAFlnk), transcription-associated (Tx, TxWk), enhancer-associated (Enh, EnhG), Polycomb-repressed (ReprPC, ReprPCWk), and quiescent states. Enrichment was summarized as log_2_ odds ratios, with brain and non-brain tissues presented separately to evaluate whether TE-derived regulatory enrichment was specific to brain chromatin context.

### 2.8 Integration of Brain Cell-Type-Specific Chromatin Accessibility

To characterize the regulatory context of TE-embedded GWAS variants, we integrated chromatin accessibility data from single-nucleus ATAC-seq (snATAC-seq) profiling of human brain tissue [10]. Publicly available peak sets representing major brain cell populations, including excitatory neurons, inhibitory neurons, astrocytes, oligodendrocytes, oligodendrocyte precursor cells (OPCs), microglia, and vascular-associated cells, were used to represent cell-type-specific accessible chromatin landscapes. GWAS variants were intersected with cell-type-specific ATAC peaks using BEDTools intersect, with overlaps defined as ≥1 bp. Chromatin accessibility was treated as a binary feature (peak present or absent), and no normalization of ATAC signal intensity was performed because analyses focused on enrichment of variant localization rather than quantitative accessibility differences.

### 2.9 Motif Enrichment Analysis of TE-Associated Regulatory Elements

To further investigate whether TE-overlapping regulatory regions harbor distinct transcription factor (TF) binding architectures, we performed sequence motif enrichment analysis using HOMER (v4.10) [6]. Human candidate cis-regulatory elements (cCREs) were obtained from the ENCODE registry (GRCh38-cCREs) as mentioned above. Foreground regions were defined as cCRE intervals overlapping both a TE annotation of interest and at least one brain-aging GWAS variant. Background regions consisted of cCRE intervals overlapping the same TE annotation but lacking GWAS variant overlap. This matched-background strategy was used to control for baseline sequence composition and regulatory context associated with each TE family. For computational efficiency and balanced comparison, background regions were randomly downsampled to approximately tenfold the number of foreground regions.

Motif enrichment was performed using findMotifsGenome.pl against the hg38 reference genome with the exact genomic interval sizes retained (-size given). Analyses included both known motif scanning and de novo motif discovery using motif lengths of 8, 10, and 12 base pairs. Parallel computation was enabled where available.

Initial analyses were conducted for Alu-overlapping cCREs and expanded to all SINE-overlapping cCREs to assess whether motif patterns were specific to the Alu subset or generalized across the broader SINE class. Enriched motifs were ranked by hypergeometric significance, with multiple-testing correction reported using the Benjamini–Hochberg false discovery rate. Resulting motifs were subsequently grouped into broader TF families (e.g., ETS/IRF, nuclear receptor, zinc finger, homeobox) to facilitate biological interpretation.

### 2.10 Activity-by-Contact (ABC) enhancer–gene prediction analysis

To identify target genes of TE-derived regulatory variants, we integrated brain-aging GWAS variants overlapping TE-derived cCREs with published Activity-by-Contact (ABC) enhancer–gene predictions [4, 13]. ABC predictions were filtered using the published ABC score threshold of 0.015 and converted to BED format using enhancer coordinates, target gene, ABC score, and biosample annotation. Because enhancer predictions were available across multiple biosamples, duplicate SNP–gene associations were collapsed to unique SNP–gene pairs while retaining biosample information for downstream tissue-specific interpretation. TE-derived cCRE-overlapping GWAS variants were intersected with ABC enhancer intervals using BEDTools. Unique SNP–gene–biosample links were retained, and target genes were summarized across all overlapping variants.

Gene Ontology enrichment analysis of ABC-predicted target genes was performed using the Database for Annotation, Visualization, and Integrated Discovery (DAVID) v2025q3 [17], querying the Gene Ontology Biological Process, Molecular Function, and Cellular Component categories together with the KEGG Pathway database, using *Homo sapiens* as the background population. Terms were considered significantly enriched at Benjamini–Hochberg FDR < 0.05. For locus-specific interpretation, we prioritized TE-derived regulatory variants with ABC links to genes with established relevance to brain aging or neurodegeneration. Linkage disequilibrium between variants at highlighted loci was calculated using PLINK and the 1000 Genomes European reference panel.

### 2.11 Experimental Enhancer–Promoter Interaction Analysis

To investigate potential downstream regulatory targets of TE-derived regulatory elements, we integrated experimentally derived enhancer–promoter and promoter–promoter interaction data from human neural progenitor cells (hNPCs), obtained via RIC-seq [8]. The source dataset reported one row per detected interaction, including the identity and genomic coordinates of both the enhancer/promoter anchor and its interacting promoter partner, an interaction p-value, and an interaction-type label (enhancer–promoter [E–P] or promoter–promoter [P–P]). Analyses were restricted to E–P interactions. Anchor coordinates were originally reported in the hg19 reference genome. To preserve the full interaction-level annotation (partner promoter identity and genomic location) through coordinate conversion, anchor coordinates were extracted, converted to hg38 using UCSC LiftOver, and rejoined to the original interaction table using each interaction’s unique identifier. Interactions whose anchor failed to lift over were excluded (30 of 52,173 interactions; 0.06%).

Brain-aging GWAS variants overlapping TE-derived cCREs were intersected with hg38-lifted hNPC enhancer/promoter anchors using BEDTools intersect. Because a single anchor region can participate in multiple independent interactions with different promoter partners, the resulting overlaps were summarized at two levels: (1) the number of unique anchor regions overlapping a TE-derived cCRE-associated GWAS variant, and (2) for each such anchor, the number of unique promoter partners (distinct interaction targets) linked to it across all retained E–P interactions. This distinct-promoter count was used as the primary measure of an anchor’s regulatory connectivity. Resulting enhancer–promoter pairs were used to identify candidate target genes associated with TE-derived regulatory loci and to characterize the potential downstream regulatory networks connected to TE-derived GWAS signals.

### 2.12 Multiome-based Brain Aging Genetic ScoreCard for prioritization of TE-embedded regulatory variants

To prioritize candidate functional regulatory variants, we developed a multiomic evidence-integration framework that combined orthogonal regulatory annotations and variant-to-gene (V2G) evidence generated throughout this study. Analysis was restricted to independent genomic loci, defined by linkage disequilibrium (LD)-based clumping of genome-wide significant variants (PLINK v1.9; *r*^2^ > 0.8, 5 Mb clumping window, 1000 Genomes Phase 3 European reference panel), yielding 38 independent loci genome-wide. For each locus, TE-derived cCRE-overlapping GWAS variants were evaluated across 27 independent evidence layers, and the locus was represented by its highest-scoring variant, preventing loci with disproportionately large numbers of candidate variants from being over-represented in downstream prioritization. The complete set of 633 TE-derived cCRE-overlapping variants, scored individually prior to locus-level collapsing, is provided in Supplementary Table 2. This approach parallels multiomic scorecard frameworks recently applied to prioritize convergent coding and regulatory variants in other complex trait contexts, such as kidney function [9].

Evidence was organized into two components. A *variant-level regulatory score* (maximum = 8) captured evidence intrinsic to the variant’s own regulatory context, independent of any specific target gene: TE-derived cCRE membership, enhancer-like cCRE annotation (dELS or pELS), membership in a TE family of interest (Alu, SVA, TcMar-Tigger, or MIR), proximity to a TE-derived transcription start site, overlap with a regulatory ChromHMM state, overlap with an hNPC enhancer–promoter interaction anchor, presence of any Activity-by-Contact (ABC) enhancer–gene link, and presence of a neural-tissue-specific ABC link.

A *target-gene score* (maximum = 19) was compiled from variant-to-gene (V2G) evidence, comprising brain-specific molecular QTL datasets (eQTL, sQTL, apaQTL, fine-mapped eQTL, fine-mapped sQTL, and fine-mapped apaQTL), chromatin interaction resources (Hi-C, HiChIP, and promoter capture Hi-C), enhancer–gene linking methods (ENCODE-rE2G, EpiMap, Roadmap, and GPMap colocalization), chromatin accessibility QTLs (haQTL and single-cell eQTLs), ABC enhancer–gene links, and genome-wide integrative SNP-to-gene resources (cS2G, Open Targets Genetics, and GeneHancer). Because a single variant can be linked to multiple candidate genes with unequal evidentiary support, target-gene evidence was evaluated separately for each candidate gene, and each variant was assigned to the single gene supported by the largest number of independent V2G sources, while evidence for competing candidate genes was not pooled or attributed to the selected target gene. Gene symbols were resolved from Ensembl gene identifiers using the GENCODE v44 annotation, and each variant’s position relative to its assigned target gene (intergenic, intronic, exonic, coding, or untranslated region) was classified using the same annotation.

For each evidence layer, variants received a binary score indicating the presence or absence of supporting evidence. The variant-level and target-gene scores were summed to generate a composite prioritization score (maximum = 27), with higher scores reflecting support from a greater number of independent regulatory and V2G resources. This framework was used to rank TE-derived regulatory variants and their assigned target genes for downstream functional characterization. The complete list of evidence layers, source datasets, brain tissues, and references is provided in Supplementary Table S1. All genome-wide significant variants were retained because the objective was candidate variant prioritization within established loci rather than statistical enrichment testing.

## 3 Results

### 3.1 Brain-aging GWAS variants localize within TE-derived regulatory elements, concentrated in specific TE subclasses

We first characterized how brain-aging–associated variants are distributed across TE-derived regulatory elements. Treating the 6,323 genome-wide significant variants as candidate causal variants within the 38 associated loci (Methods), we intersected them with TE-derived candidate cis-regulatory elements (cCREs), defined as cCREs for which at least 50% of the element overlapped a single TE annotation. Of the 6,323 candidate variants, 633 (10.01%) overlapped a TE-derived cCRE, corresponding to 367 distinct TE-derived cCREs (Figure 1A). These elements were predominantly distal enhancer-like signatures (dELS, 60.5%), followed by transcription-factor elements (TF, 12.3%), chromatin-accessible elements (CA, 10.4%) and proximal enhancer-like signatures (pELS, 7.9%). As an independent validation, we integrated the candidate variants with the catalog of approximately 26,056 TE-derived transcription start sites (TE-TSSs). By expanding each TE-TSS by ±500 bp to capture promoter-proximal variation, 81 candidate variants localized within TE-derived promoter regions. These results indicate that brain-aging candidate variants overlapping TE-derived cCREs fall overwhelmingly within enhancer-type regulatory annotations.

**Figure 1:**
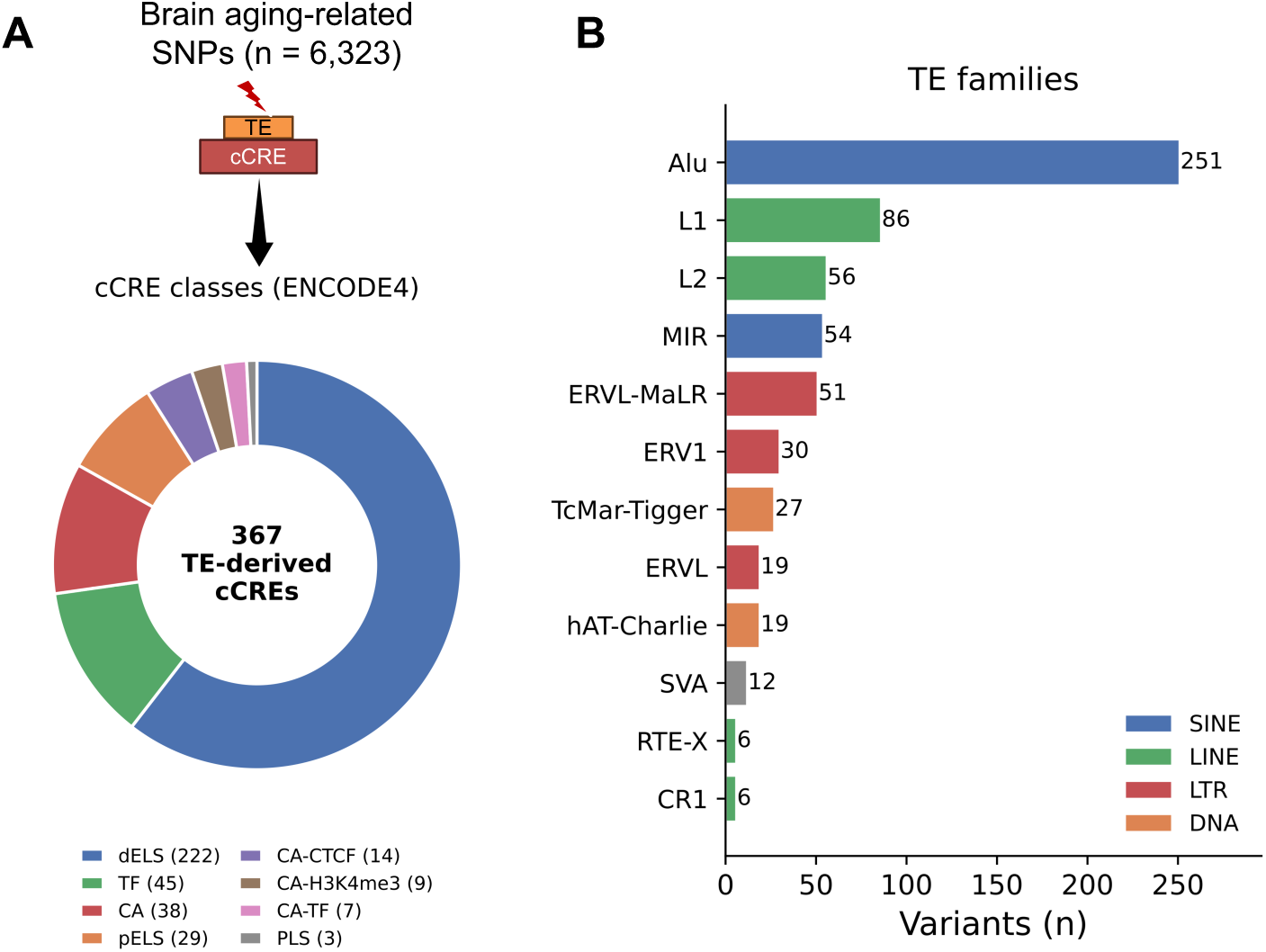
Localization and composition of brain-aging GWAS variants within TE-derived regulatory elements (A) cCRE class composition of GWAS variants overlapping TE-derived regulatory elements. (B) Number of GWAS variants overlapping TE families.

Next, we examined which TE families contributed these regulatory elements. Among the 633 TE-cCRE-overlapping candidate variants, Alu elements were the largest contributor (39.7%), followed by L1 (13.6%), L2 (8.8%), MIR (8.5%), ERVL-MaLR (8.1%), ERV1 (4.7%) and TcMar-Tigger (4.3%) (Figure ??B). This composition is dominated by SINE-derived elements (Alu and MIR together, 48.2%), consistent with the high genomic copy number of these families (Figure 1B). Together, these results show that brain-aging candidate variants are distributed across TE-derived regulatory elements, predominantly Alu and MIR-derived distal enhancers, providing a set of candidate regulatory variants for downstream prioritization.

### 3.2 TE-derived regulatory variants occupy distinct, brain-enriched chromatin-state environments

To examine tissue-specific regulatory context, we integrated TE-derived cCRE-associated GWAS variants with Roadmap Epigenomics ChromHMM annotations across six brain tissues (hippocampus, dorsolateral prefrontal cortex, substantia nigra, angular gyrus, caudate, and cingulate gyrus) and two non-brain comparator tissues (liver and fetal kidney). Enrichment was evaluated relative to all TE-derived cCREs to determine whether GWAS-associated regulatory elements preferentially occupy specific chromatin states, and whether this pattern was specific to brain tissue.

Across all six brain tissues, TE-derived cCRE-associated GWAS variants exhibited highly consistent chromatin-state profiles (Figure 2A). The strongest positive enrichment was observed for active enhancer states (Enh), with odds ratios ranging from 5.63 in hippocampus to 6.52 in angular gyrus (all FDR < 10^−56^), together with weaker enrichment for genic enhancers (EnhG) and transcription-associated states (TxWk). In contrast, quiescent chromatin showed the strongest and most consistent depletion across every tissue, accompanied by substantial depletion of heterochromatin, indicating that TE-derived regulatory variants are preferentially localized within accessible regulatory environments rather than inactive genomic regions.

**Figure 2:**
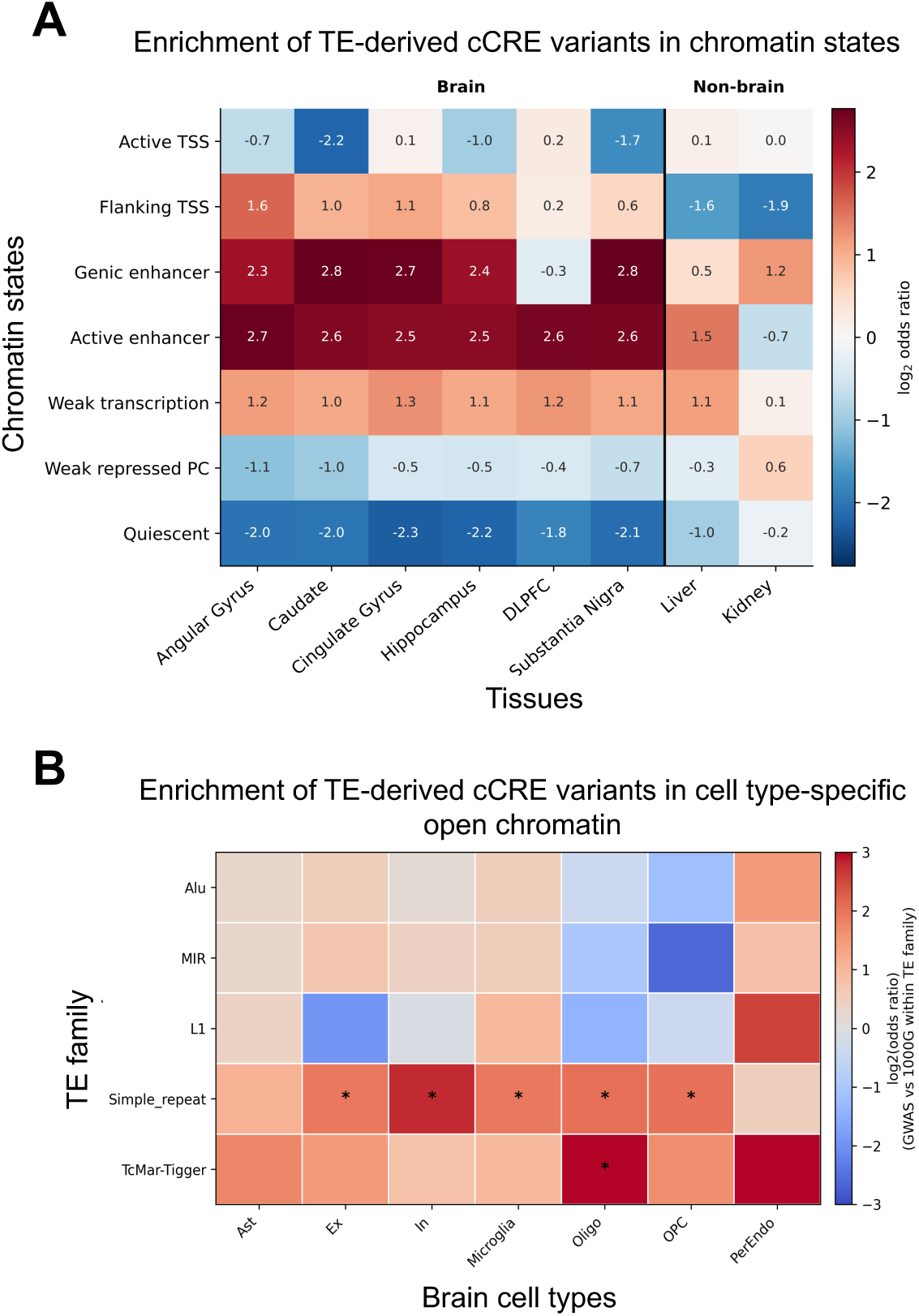
Enrichment of TE-derived cCRE-associated GWAS variants across chromatin states in brain regions and cell types. (A) Enrichment of TE-derived cCRE-associated GWAS variants across chromatin states defined by histone modifications across brain regions. (B) Cell-type-specific chromatin accessibility enrichment of brain-aging GWAS variants overlapping TE families across major brain cell populations.

To assess whether this enhancer enrichment was specific to brain tissue, we repeated this analysis in liver and kidney. Active enhancer enrichment was substantially weaker in both non-brain tissues than in any of the six brain tissues. Kidney showed no significant enrichment (OR = 0.63, FDR = 0.23), while liver showed diminished but still nominally significant enrichment (OR = 2.75, FDR = 1.4 × 10^−19^), roughly half the effect size observed in brain tissue. This pattern indicates that active-enhancer enrichment among TE-derived cCRE-associated GWAS variants is substantially stronger and more consistent in brain tissue than in non-brain tissue, though not exclusively restricted to brain, consistent with a brain-enriched rather than strictly brain-specific regulatory signature.

Weak Polycomb-repressed chromatin (ReprPCWk) also demonstrated reproducible enrichment across multiple brain regions. Although these loci are not uniformly associated with fully active regulatory states, their enrichment within Polycomb-associated chromatin suggests that a subset of TE-derived regulatory elements may reside in developmentally poised or context-dependent regulatory environments that retain the potential for transcriptional activation.

Overall, the chromatin-state enrichment profiles were highly reproducible across all six brain tissues and clearly distinguishable from the weaker or absent enrichment observed in non-brain tissue, supporting the conclusion that TE-derived regulatory variants associated with brain aging preferentially occupy accessible, brain-enriched regulatory chromatin environments rather than being randomly distributed throughout the genome or non-specifically active across all tissue types.

### 3.3 TE-overlapping GWAS variants show cell-type-specific chromatin accessibility patterns

To evaluate whether TE-overlapping GWAS variants were preferentially located in accessible chromatin across brain cell types, we intersected TE-embedded variants with cell-type-specific ATAC-seq peaks from major brain cell populations, including excitatory neurons, inhibitory neurons, astrocytes, oligodendrocytes, oligodendrocyte precursor cells (OPCs), microglia, and vascular-associated cells.

This analysis revealed non-random enrichment patterns across TE families and cell types. Simple repeat elements showed the most consistent enrichment across multiple cell types, including excitatory neurons (OR = 3.83, FDR = 0.0017), inhibitory neurons (OR = 6.82, FDR = 4.0 × 10^−6^), microglia (OR = 3.79, FDR = 0.0339), OPCs (OR = 4.07, FDR = 0.0126), and oligodendrocytes (OR = 4.12, FDR = 0.0058). Oligodendrocytes also showed strong enrichment for TcMar-Tigger elements (OR = 11.84, FDR = 0.0026), representing the largest cell-type-specific effect observed (Figure 2B). These findings suggest that regulatory accessibility among TE-overlapping brain-aging variants is concentrated in specific repeat families and cellular contexts rather than broadly distributed across all TE classes.

### 3.4 TE-overlapping variants are enriched in transcription factor-associated regulatory contexts

We next asked whether TE-overlapping GWAS variants differed from non-TE variants in their cCRE regulatory architecture. Distal enhancer-like signatures were the most common cCRE class among both TE-overlapping and non-TE GWAS variants, consistent with the enhancer-rich architecture of noncoding GWAS signals. However, TE-overlapping variants showed a distinct regulatory distribution, with increased representation in transcription factor-associated regulatory classes and relative depletion from promoter-like signatures (Figure 3A).

**Figure 3:**
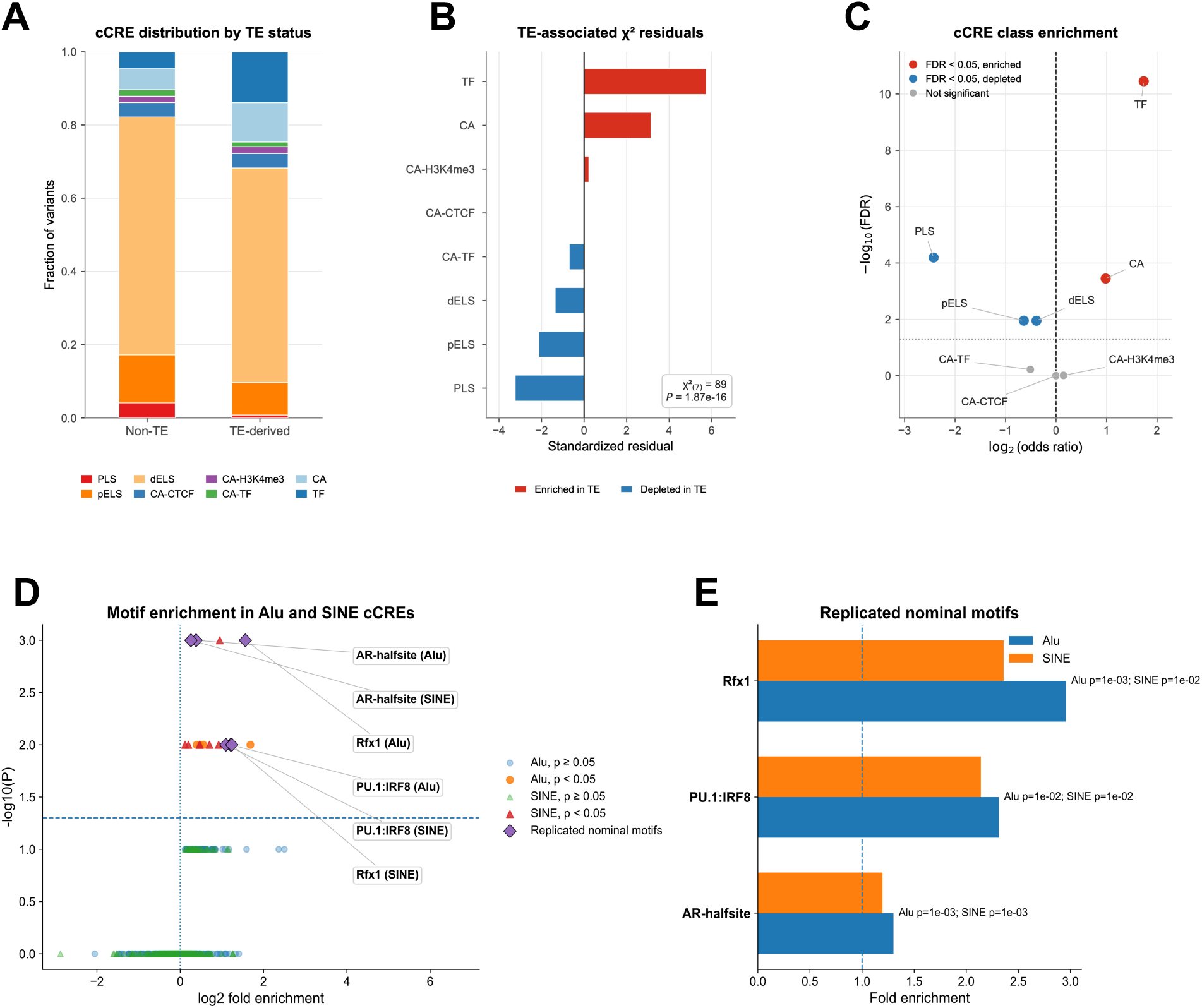
Regulatory architecture of TE-overlapping and non-TE GWAS variants. (A) cCRE class distribution by TE-overlap status. (B) Standardized residuals from chi-square analysis showing cCRE classes contributing to TE-associated regulatory differences. (C) cCRE class enrichment among TE-overlapping variants. (D) Combined motif enrichment volcano plot for Alu and SINE analyses. (E) Nominally replicated motifs observed in both Alu and SINE analyses.

A chi-square test confirmed a significant association between TE-overlap status and cCRE class distribution (*p* = 2.21 × 10^−25^). Standardized residual analysis showed that transcription factor-associated cCREs contributed strongly to the TE-associated signal, with CA-TF and CA-H3K4me3 classes also overrepresented among TE-overlapping variants. In contrast, promoter-like signatures showed negative residuals, indicating depletion among TE-overlapping variants (Figure 3B). Fisher’s exact tests across individual cCRE classes further supported this pattern, showing enrichment of transcription factor-associated regulatory classes and depletion of promoter-like elements among TE-overlapping variants (Figure 3C). These results indicate that TE-overlapping GWAS variants are preferentially embedded in transcription factor-associated regulatory environments rather than core promoter regions.

### 3.5 Motif enrichment reveals modest and heterogeneous regulatory signatures in SINE-associated cCREs

Because SINE elements, in particular Alu, showed evidence of enrichment among brain-aging GWAS loci, we next tested whether SINE-associated regulatory elements shared coherent transcription factor motif architecture. HOMER motif enrichment analysis was performed separately for Alu-overlapping cCREs and for all SINE-overlapping cCREs, using TE-matched cCRE backgrounds without GWAS overlap.

For Alu-associated regions, 290 foreground cCREs were compared against matched Alu-overlapping background regions. Several motifs showed nominal enrichment, including AR-halfsite (FE = 1.30, *p* = 10^−3^), Rfx1 (FE = 2.96, *p* = 10^−3^), and PU.1 (FE = 2.31, *p* = 10^−2^). Expanding the analysis to all SINE-overlapping cCREs increased the foreground set to 415 regulatory regions and yielded a similar but heterogeneous pattern, with nominal enrichment of AR-halfsite, Rfx1, PU.1, and additional nuclear receptor, immune-associated, and developmental motifs (Figure 3D).

Three motifs, AR-halfsite, PU.1, and Rfx1, were nominally enriched in both Alu-specific and broader SINE analyses (Figure 3E). However, no motif remained significant after multiple-testing correction. These results suggest that SINE-associated regulatory loci do not converge on a single dominant motif program. Instead, motif architecture appears modest and heterogeneous, with recurrent but nominal signatures related to nuclear receptor signaling, immune-associated transcriptional regulation, and chromatin-associated transcription factors.

### 3.6 Enhancer–gene predictions link TE-derived regulatory variants to candidate target genes

To identify potential downstream targets of TE-derived regulatory variants, we intersected TE-derived cCRE-overlapping GWAS variants with Activity-by-Contact (ABC) enhancer–gene predictions [4, 13]. Across all biosamples, this analysis identified 529 enhancer–gene links involving 79 GWAS variants and 89 unique target genes. Gene Ontology enrichment analysis of ABC-predicted target genes did not identify any significantly enriched terms after multiple-testing correction (all Benjamini-adjusted *P* = 1.0), indicating that ABC-linked target genes are not concentrated within a specific functional pathway.

Several TE-derived regulatory elements exhibited extensive predicted enhancer–gene connectivity, linking individual cCREs to multiple protein-coding genes, long noncoding RNAs, and regulatory transcripts. This network organization suggests that TE-derived regulatory elements may influence coordinated transcriptional programs rather than single target genes. The most prominent example occurred within the chromosome 17q21.31 inversion locus, where a MIR-derived distal enhancer containing multiple highly linked GWAS variants was repeatedly connected by the ABC model to *MAPT*, *MAPT-AS1*, and *MAPT-IT1*, with *TRIM47* and *SPATS2L* among the most extensively connected target genes genome-wide (Figure 4A). Together, these findings identify TE-derived regulatory elements as candidate hubs capable of integrating disease-associated genetic variation with complex enhancer–promoter interaction networks, independent of any single overrepresented biological pathway.

**Figure 4:**
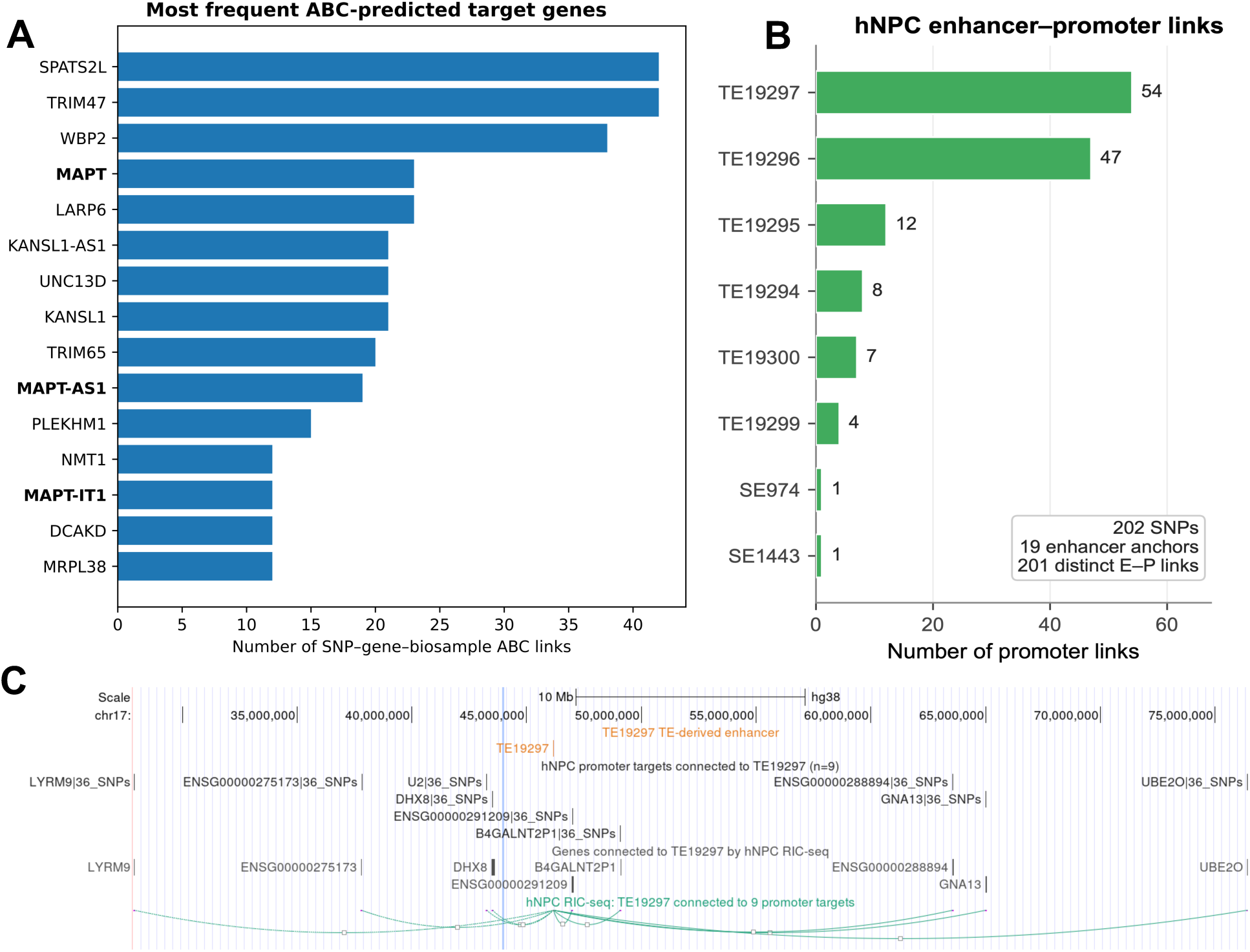
(A) Most frequently observed ABC target genes across TE-derived cCRE-overlapping GWAS variants. (B) hNPC enhancer anchors overlapping TE-derived cCRE GWAS variants and their promoter interaction counts. (C) Genome browser view of TE19297, the most extensively connected TE-derived enhancer anchor, showing its complete experimentally supported hNPC RIC-seq connectivity to promoter targets.

### 3.7 Neural enhancer–promoter interactions support TE-derived regulatory elements as candidate gene regulators

To determine whether TE-derived regulatory elements participate in experimentally observed neural regulatory interactions, we intersected TE-derived cCREs containing brain-aging GWAS variants with enhancer anchors from human neural progenitor cell (hNPC) enhancer–promoter interaction maps, using the complete RIC-seq interaction-level dataset to preserve individual promoter identity and genomic location for each interaction. This analysis identified 202 GWAS variants located within 19 TE-derived enhancer anchors that collectively formed 201 experimentally supported enhancer–promoter interactions, reaching 192 distinct promoter targets (Figure 4B), with the complete network shown in Supplementary Figure S1. These enhancer anchors connected to numerous protein-coding genes and regulatory transcripts, demonstrating that TE-derived cCREs participate in extensive regulatory interaction networks within a neural developmental context.

Several enhancer anchors exhibited particularly broad regulatory connectivity, with the most highly connected elements concentrated at the chromosome 17q21.31 locus, as one enhancer, TE19297, anchored 36 GWAS variants and formed interactions with 54 promoter regions, while a neighboring anchor connected 52 variants to 47 promoter regions. These observations suggest that a subset of TE-derived regulatory elements functions as highly connected regulatory hubs capable of coordinating transcription across multiple genes. To illustrate this connectivity directly, Figure 4C shows the complete experimentally supported hNPC RIC-seq connectivity of TE19297, the most extensively connected TE-derived enhancer anchor identified in this analysis.

### 3.8 Brain Aging Genetic ScoreCard prioritizes TE-derived regulatory elements for functional follow-up

To prioritize candidate brain aging loci, we integrated regulatory annotations, chromatin state information, and variant-to-gene evidence into a multiomic prioritization framework, the Brain Aging Genetic ScoreCard (Figure 5). Analysis was restricted to independent genomic loci, defined by LD-based clumping (*r*^2^ > 0.8), and each locus was represented by its highest-scoring TE-derived regulatory variant. Across 27 orthogonal evidence layers, the Brain Aging Genetic ScoreCard identified 25 independent loci with a TE-derived candidate regulatory variant and an assigned target gene. Using prioritization score of at least 10 (supported by ≥10 independent evidence layers), we identified 10 high-confidence brain-aging-related genes, including MAPT, TRIM47, SLC16A8, LARP6, SPATS2L, ZFAND2A, HLA-C, PLD1, ICA1L and EIF3M (Figure 5A).

**Figure 5:**
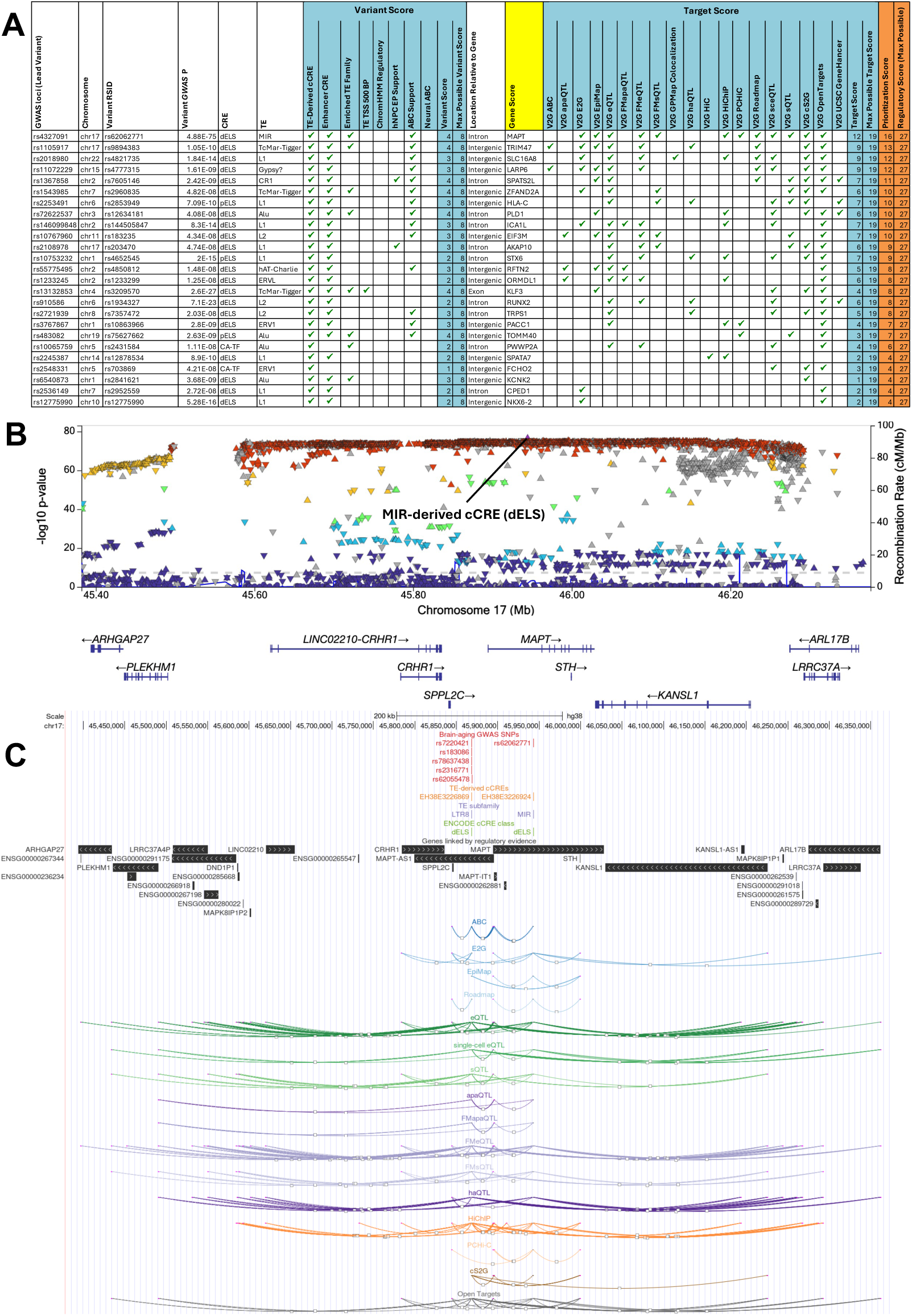
Brain Aging Genetic ScoreCard prioritizes a MIR-derived regulatory element at the chromosome 17q21.31 MAPT locus. (A) Multiomic scorecard summarizing regulatory and variant-to-gene evidence across 27 orthogonal layers for the top tne highest-priority TE-derived variants. (B) Regional association plot showing the GWAS signal with LD (r²) relative to the prioritized variant. (C) Genome browser view of enhancer–gene and variant-to-gene evidence, integrating ABC predictions, hNPC enhancer–promoter interactions, and molecular QTL, chromatin interaction, and integrative V2G resources.

Among these, the chromosome 17q21.31 *MAPT* locus emerged as the highest-confidence candidate, with a prioritization score of 16, the highest of any locus genome-wide and substantially above the next-highest locus (score of 13). A MIR-derived distal enhancer-like cCRE containing multiple highly associated GWAS variants accumulated support from enhancer annotation, TE-derived promoter proximity, ABC enhancer predictions, brain ChromHMM states, and numerous independent variant-to-gene resources (Figure 5A). Because raw evidence accumulation could in principle scale with the number of candidate variants per locus, we assessed whether this ranking was influenced by locus size. Modest, statistically significant residual correlations remained between locus size and both the variant-level regulatory score (Spearman *ρ* = 0.35, *P* = 0.083, not significant) and the target-gene evidence score (*p* = 0.51, *P* = 0.008) after restricting scoring to one representative variant per locus, substantially attenuated relative to an unadjusted, per-variant scoring scheme evaluated during method development (*p* ≈ 1.0). Critically, *MAPT*’s top rank was robust to this residual confound, its prioritization score was not disproportionate relative to substantially smaller loci despite containing more genome-wide significant variants (n = 802, within an extended 900-kb 17q21.31 inversion haplotype) than any other locus. For example, a locus with only 11 candidate variants (*TRIM47*) scored 13, closely approaching *MAPT*’s score of 16. We therefore report *MAPT*’s prioritization as reflecting convergent, independently derived regulatory evidence at a specific enhancer element on a confound-resistant scoring framework, while noting that a modest, quantified relationship between locus size and target-gene evidence remains and should be considered when interpreting relative rankings among lower-scoring loci.

We selected the *MAPT* locus for detailed functional characterization. Regional association analysis demonstrated that the prioritized MIR-derived enhancer resides within the peak of a strong brain-aging GWAS association signal (Figure 5B). Linkage disequilibrium analysis using the 1000 Genomes European reference panel showed that the prioritized variants span multiple LD structures within the 17q21.31 locus, as three prioritized variants (rs78637438, rs2316771, and rs62055478) formed a complete LD block (*r*^2^ = 1.0) whereas rs7220421 and rs183086 exhibited only weak to moderate correlation with this block (*r*^2^ = 0.18 − 0.36). These findings indicate that the Brain Aging Genetic ScoreCard prioritizes several regulatory elements across the broader *MAPT* locus rather than a single perfectly linked association signal. Because these variants reside within the extended 17q21.31 inversion haplotype, their individual regulatory contributions cannot currently be distinguished from broader haplotype-level effects.

ABC enhancer–gene predictions linked the prioritized TE-derived regulatory element to *MAPT*, *MAPT-AS1*, *MAPT-IT1*, and *SPPL2C* across 16 SNP-gene links. Considering all variant-to-gene evidence together, *MAPT* was the most consistently supported target gene at this element, with independent support from 13 of 16 evidence sources with any data at this locus (ABC, E2G, EpiMap, HiChIP, promoter capture Hi-C, Open Targets, Roadmap, cS2G, eQTL, sQTL, single-cell eQTL, and both fine-mapped eQTL and sQTL). Promoter capture Hi-C support was derived from a single prioritized variant and should be interpreted as preliminary. This regulatory element’s connectivity was not exclusive to *MAPT*, as several neighboring genes, including *PLEKHM1*, *SPPL2C*, *KANSL1*, *MAPT-AS1*, *LRRC37A*, *CRHR1*, and *STH*, also received substantial independent support (7–9 of 16 sources each), consistent with the extended 17q21.31 locus containing multiple co-regulated genes (Figure 5C; complete variant-level evidence for all TE-derived cCREs in this and other loci is provided in Supplementary Table 2).

Together with the multiomic evidence summarized in the Brain Aging Genetic ScoreCard, these findings identify this TE-derived enhancer as a strong candidate regulatory element influencing expression of genes within the *MAPT* locus. Dysregulation of *MAPT* expression promotes the accumulation of pathological tau aggregates, a hallmark of tauopathies including Alzheimer’s disease and frontotemporal dementia [18]. To our knowledge, this is the first study to implicate a TE-derived regulatory element in *MAPT* transcriptional regulation specifically in the context of brain age gap. Collectively, these results illustrate how integrative functional genomics can prioritize TE-derived regulatory elements that likely mediate the effects of noncoding GWAS variation.

## 4 Discussion

In this study, we developed an integrative genomic framework to investigate how brain-aging GWAS variants intersect transposable element-derived regulatory architecture. Rather than treating all TE-overlapping variants as a single category, we distinguished broad TE sequence overlap from experimentally annotated TE-derived regulatory elements, cell-type-specific chromatin accessibility, cCRE regulatory class, enhancer–promoter connectivity, motif architecture, and SNP-based heritability contribution. Our results support a model in which a subset of noncoding brain-aging variants localize to TE-derived regulatory elements that can be prioritized using integrated functional genomic evidence.

A central observation is the localization of brain-aging–associated variants within TE-derived regulatory elements, including promoters and distal enhancers. Of the 6,323 genome-wide-significant variants, 633 (10.0%) localized within 367 TE-derived cCREs, most of which were enhancer-like. Examining the composition of this TE-derived variant set, Alu-derived elements formed the largest family component, followed by L1, L2, MIR and ERVL-MaLR elements. Further, analysis of tissue-specific chromatin states and cell-type-specific chromatin accessibility indicated that this TE-associated regulatory architecture is context-dependent. Across brain cell types, accessible TE-derived elements were distributed non-uniformly among repeat families and cellular contexts. Simple-repeat elements were accessible across multiple brain cell types, whereas TcMar-Tigger elements showed a predominantly oligodendrocyte signal. These observations suggest that different repeat families may contribute to brain-aging GWAS interpretation through distinct cellular regulatory contexts. The oligodendrocyte signal is notable given the importance of myelination, white-matter integrity and glial regulation in brain aging, although our analysis remains correlative and does not establish that TcMar-Tigger elements causally affect oligodendrocyte function.

Consistent with the predominance of enhancer-like TE-derived cCREs, our analyses indicate that TE-derived cCRE-overlapping variants are embedded within transcription factor–associated and open-chromatin regulatory environments rather than promoter-centric architecture. Among TE-derived cCREs, transcription factor–associated (TF) elements were the most common class relative to non-TE cCREs, while promoter-like signatures were comparatively rare. Motif analysis of the full brain-aging GWAS provided independent, sequence-level support for this interpretation, as several motifs were nominally and reproducibly enriched across Alu and broader SINE analyses, including AR-halfsite, PU.1:IRF8, and RFX-family motifs. These signals point to possible involvement of nuclear-receptor signaling, immune-associated transcriptional programs, and chromatin-regulatory pathways. PU.1:IRF8 is of particular interest in the context of microglial and immune regulation, whereas RFX-family factors are relevant to chromatin accessibility and neural gene regulation.

Finally, we developed the Brain Aging Genetic ScoreCard, which integrates 27 orthogonal regulatory evidence layers, including enhancer annotations, molecular QTLs, chromatin interaction maps, enhancer– promoter predictions, and integrated variant-to-gene resources, evaluated at the level of independent, LD-clumped genomic loci to avoid conflating locus size with regulatory support. Applying this framework identified 25 independent loci with a TE-derived candidate regulatory variant and prioritized 10 high-confidence brain-aging-associated target genes, including *MAPT*, which reflects convergent support from enhancer annotation, TE-derived promoter proximity, ABC enhancer predictions, brain ChromHMM states, and numerous independent variant-to-gene resources at a specific MIR-derived enhancer.

Several limitations should be considered when interpreting these results. First, the map of TE-derived cCRE variants associated with brain aging is still incomplete, which may be expanded with further GWAS studies in larger sample sizes from different ancestries. Second, the TE-derived cCRE atlas used in this study was derived from a broad collection of ENCODE epigenomic datasets rather than aging-specific brain tissue. To minimize this bias, we incorporated brain-specific chromatin states and cell-type-specific chromatin accessibility, which resolved brain cell-type-specific regulation among brain-aging-related variants. Third, the single-nucleus ATAC-seq reference used for cell-type-specific accessibility analyses was derived from a case–control cohort profiled for Alzheimer’s disease rather than from a cohort specifically designed to characterize normative brain aging. Future work incorporating chromatin-accessibility atlases derived from cognitively healthy, age-diverse cohorts specifically profiled for aging-related regulatory change would help disambiguate disease-specific from generalizable aging-related regulatory signal. Last, this study is mainly focused on brain aging due to data availability. Further expansion to other organs will be essential to uncover the functions of TE-derived cCRE variants in human aging and age-related diseases.

Despite these limitations, this study provides an integrative framework combining transposable element biology with regulatory genomics to interpret noncoding GWAS variation. By combining GWAS variants, TE-derived regulatory annotations, chromatin accessibility, motif analyses, enhancer–promoter interactions, and multiomic prioritization, we demonstrate that this integration can identify biologically plausible regulatory mechanisms underlying noncoding GWAS loci. Together, these findings suggest that a subset of brain-aging loci converge on TE-derived regulatory elements with independent functional support, illustrating how transposable element biology can refine the interpretation of noncoding GWAS signals.

## Supporting information

Supplemental Table Scorecard

## Data & Code Availability

All datasets are publicly available from their respective primary sources. All analysis scripts, processed data annotations, and figure-generation code will be publicly available following publication of the manuscript.

## Acknowledgement

The work in Liu’s laboratory has been supported by the NIH National Institute on Aging (grant nos. P01AG047200 and P30AG092746) and startup funding from the University of Rochester. We acknowledge the public multiomic datasets, including the brain aging GWAS and functional annotations obtained from UCSC, ENCODE, Roadmap, and other studies.

## Supplementary Figures and Tables

**Figure S1:**
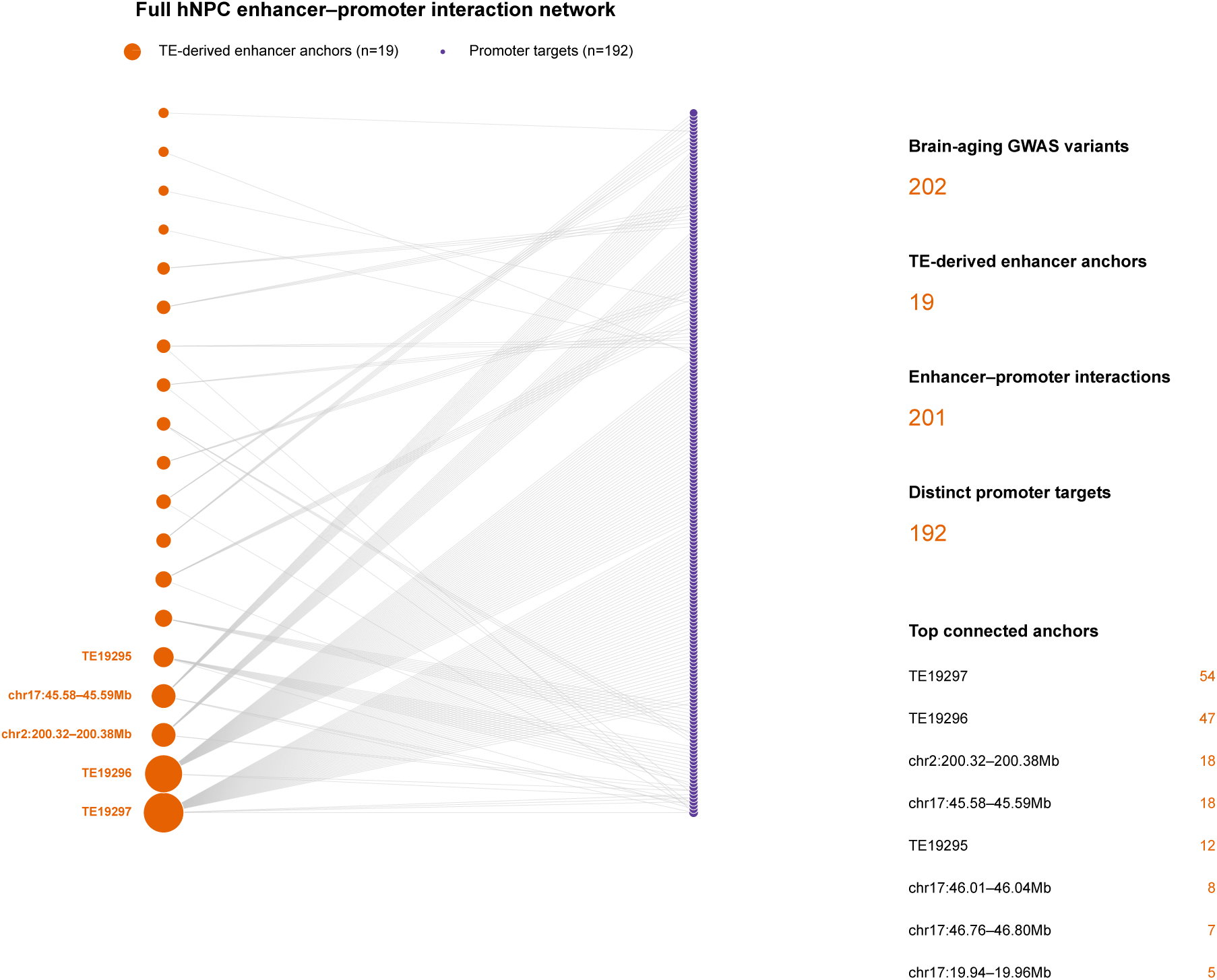
Full hNPC enhancer-promoter interaction network linking brain-aging TE-derived cCRE SNPs to regulatory targets.

**Table S1:** Variant-to-gene (V2G) resources integrated into the multiomic prioritization framework. Brain-specific datasets were used whenever available.

| Category | Resource | Brain tissues | Biological evidence |
| --- | --- | --- | --- |
| Enhancer-gene | ABC | Cortex | Activity-by-contact enhancer-gene predictions |
| Enhancer-gene | ENCODE-rE2G | Selected brain tissues | Predicted enhancer-gene regulatory links |
| Enhancer-gene | EpiMap | Selected brain tissues | Enhancer-gene interaction predictions |
| Enhancer-gene | Roadmap | Selected brain tissues | Enhancer-gene linkage predictions |
| Expression QTL | GTEx eQTL | 13 GTEx brain tissues* | Variant associations with gene expression |
| Expression QTL | Fine-mapped eQTL (SuSiE) | 13 GTEx brain tissues* | High-confidence causal eQTLs |
| Splicing QTL | GTEx sQTL | 13 GTEx brain tissues* | Variant associations with RNA splicing |
| Splicing QTL | Fine-mapped sQTL (SuSiE) | 13 GTEx brain tissues* | High-confidence causal sQTLs |
| APA QTL | GTEx apaQTL | 13 GTEx brain tissues* | Variant associations with alternative polyadenylation |
| APA QTL | Fine-mapped apaQTL (SuSiE) | 13 GTEx brain tissues* | High-confidence causal APA-QTLs |
| Chromatin interaction | Hi-C | Cortex, hippocampus | Chromatin contact maps |
| Chromatin interaction | HiChIP | Cortex | Regulatory chromatin interactions |
| Chromatin interaction | Promoter Capture Hi-C | Frontal cortex BA9, hippocampus | Promoter-centered chromatin interactions |
| Colocalization | GPMaP | 13 GTEx brain tissues* | GWAS-QTL colocalization |
| Accessibility QTL | haQTL | Cortex | Chromatin accessibility QTLs |
| Accessibility QTL | Single-cell eQTL | Cortex | Cell-type-specific expression QTLs |
| Integrated V2G | cS2G | Genome-wide | Integrated SNP-to-gene predictions |
| Integrated V2G | Open Targets Genetics | Genome-wide | Machine learning-based variant-to-gene prioritization |
| Integrated V2G | GeneHancer | Genome-wide | Curated enhancer-gene associations |

\*GTEx v11 brain tissues include amygdala, anterior cingulate cortex (BA24), caudate (basal ganglia), cerebellar hemisphere, cerebellum, cortex, frontal cortex (BA9), hippocampus, hypothalamus, nucleus accumbens (basal ganglia), putamen (basal ganglia), spinal cord (cervical C-1), and substantia nigra.

